# 24-hour multi-omics analysis of residential sewage reflects human activity and informs public health

**DOI:** 10.1101/728022

**Authors:** Mariana Matus, Claire Duvallet, Melissa Kido Soule, Sean M. Kearney, Noriko Endo, Newsha Ghaeli, Ilana Brito, Carlo Ratti, Elizabeth B. Kujawinski, Eric J. Alm

## Abstract

High-throughput molecular analysis of sewage is a promising tool for precision public health. Here, we combine sewer network and demographic data to identify a residential catchment for sampling, and explore the potential of applying untargeted genomics and metabolomics to sewage to collect actionable public health data. We find that wastewater sampled upstream in a residential catchment is representative of the human microbiome and metabolome, and we are able to identify glucuronidated compounds indicative of direct human excretion, which are typically degraded too quickly to be detected at treatment plants. We show that diurnal variations during 24-hour sampling can be leveraged to discriminate between biomarkers in sewage that are associated with human activity from those related to the environmental background. Finally, we putatively annotate a suite of human-associated metabolites, including pharmaceuticals, food metabolites, and biomarkers of human health and activity, suggesting that mining untargeted data derived from residential sewage can expand currently-used biomarkers with direct public health or policy relevance.

## Introduction

Nearly all human activity within our cities leaves a chemical or biological footprint in sewage, but very little has been done to mine this wealth of information for the public good (Daughton 2018). Wastewater-based epidemiology could have enormous benefit to public health practice, providing real-time assessments of population health and generating data to support public health and city policies, while current practice often relies on survey-based data and evaluation of indirect outcomes (Falbe 2016, Escobar 2013). Public health surveillance is primarily based on reports from hospitals and other clinical centers, and appropriate responses are initiated after a threshold of clinical cases have been reported. For example, wastewater-based public health surveillance platforms could provide data more effectively than hospital-based reporting, allowing public health officials to respond to emerging outbreaks before a critical mass of patients have gotten sick (Manor 2014, Hellmer 2014, Yan 2018). In addition to identifying infectious disease outbreaks, wastewater could provide a novel data source to evaluate the effectiveness of policies addressing long-term population health (Daughton 2018, Burgard 2014), providing a more immediate measure of the impact of health-related policies on targeted outcomes than existing approaches.

Recently, wastewater-based epidemiology was used as an early indicator of poliovirus outbreaks to target vaccination campaigns (Manor 2014, Kaliner 2015), to monitor asymptomatic virus circulation (Berchenko 2017), and as a novel source of data on illicit drug consumption in Australia (Australian Report) and over 60 European cities and towns (van Nuijs 2018, Thomas 2012, Baz-Lomba 2016). More recently, wastewater analysis has been proposed as a promising tool to tackle the severe opioid epidemic in the U.S. and Canada (Keshaviah 2016, Keshaviah 2017) as well as to monitor antimicrobial resistance (Pehrsson 2017, Initiatives 2018), obesity levels, and lifestyle biomarkers (Daughton 2018, Arnaud 2018, Newton 2015). Apart from polio surveillance and drug consumption estimation, many applications remain proof-of-concept academic projects and are not yet implemented into public health practices.

Current practices, such as sampling downstream sites like pump stations or wastewater treatment plants (WWTPs) (Manor 2014, Hellmer 2014, Australian Report, Thomas 2012, Baz-Lomba 2016) may not be effective for all applications because of degradation of chemical and biological molecules during transport. Moreover, grouping samples of different origins can complicate analysis by aggregating multiple municipalities with variable transit times to the collection site, and by mixing of residential and industrial waste. These factors create technical biases in the data and prevent accurate consumption or infection rates from being calculated from measured quantities (Thai 2014, Fahrenfeld 2016). Comparisons among WWTPs from different municipalities are further confounded by factors such as non-uniform population sizes (O’Brien 2013), variable wastewater transit times (Thai 2014), differences in sampling platforms (Ort 2010), and a lack of information on which biomarkers are stable and quantifiable across WWTPs (Arnaud 2018, Daughton 2018). Finally, composite samples from WWTPs also lack geographic specificity, reducing their usefulness to policy makers (Zuccato 2008, Castiglioni 2006, Thomas 2012).

Here, we demonstrate the utility of combining upstream residential sewage sampling with untargeted ‘omics analyses to obtain direct measures of human activity in residential populations, facilitating and expanding applications to public health. We combine demographic, sewage network, and geographic data sources to curate and select an upstream sampling site that reflects a residential population. We next generate microbiome and untargeted metabolomics data from a 24-hour sampling period and show that residential sewage represents the human microbiome and metabolome and reflects human diurnal activity. We putatively identify a variety of human-derived biomarkers, which can be extended to confirm biomarkers for use in a broad range of public health or policy evaluation applications.

## Results

### Upstream residential sewage contains more markers of human activity than sewage sampled at WWTPs

Sewage at WWTPs lacks geographic specificity, can represent populations of different orders of magnitude (e.g. 1,000 to 2 million people, Supp. Fig. 1), and contains sewage which has travelled on the order of minutes to hours, resulting in different rates of degradation and dilution of biological and chemical biomarkers (Newton 2015, Thai 2014). Residential catchments can be selected to have similar characteristics to each other, mitigating some confounding factors of WWTPs. We integrated GIS datasets with city-wide demographic and sewage network information to select a manhole serving an area with at least 80% residential land use, a population of over 5,000 people, and a maximum sewage hydraulic retention time of 60 minutes (Fig. 1A). We conducted two experiments: (1) a comparison of upstream grab samples with respective downstream samples and (2) a time series analysis of 24 hourly grab samples. We sampled sewage at the identified residential catchment (Fig. 1A) and a downstream pump station (N = 3 samples for each upstream and downstream sites, N = 1 upstream sample for each time point in the 24 hour experiment; Fig. 1B, Table 1) and performed untargeted metabolomics and 16S rDNA sequencing on the filtrate and cell fraction, respectively.

**Table 1.** Comparison of infrastructure characteristics of the Residential Catchment and Wastewater Treatment Plant (WWTP) Catchment sampled in this study. MGD = millions of gallons per day.

| **Variable** | **Residential Catchment** | **WWTP Catchment** |
| --- | --- | --- |
| Geographic resolution | 1 residential neighborhood | 18 municipalities |
| Population served | approx. 5,300 people | approx. 642,000 people |
| Land use | 84% residential | 44% residential |
| Average sewage hydraulic retention time | 25 minutes | 4 hours |
| Maximum sewage hydraulic retention time | 45 minutes | 10 hours |
| Average flow rate | 0.4 MGD (all days) | 94.4 MGD (all days)<br>81.2 MGD (dry days) |
| Sewer type | 100% Gravity-fed. | Mostly gravity-fed.<br>Several upstream subarea pump stations. |

**Figure 1.**
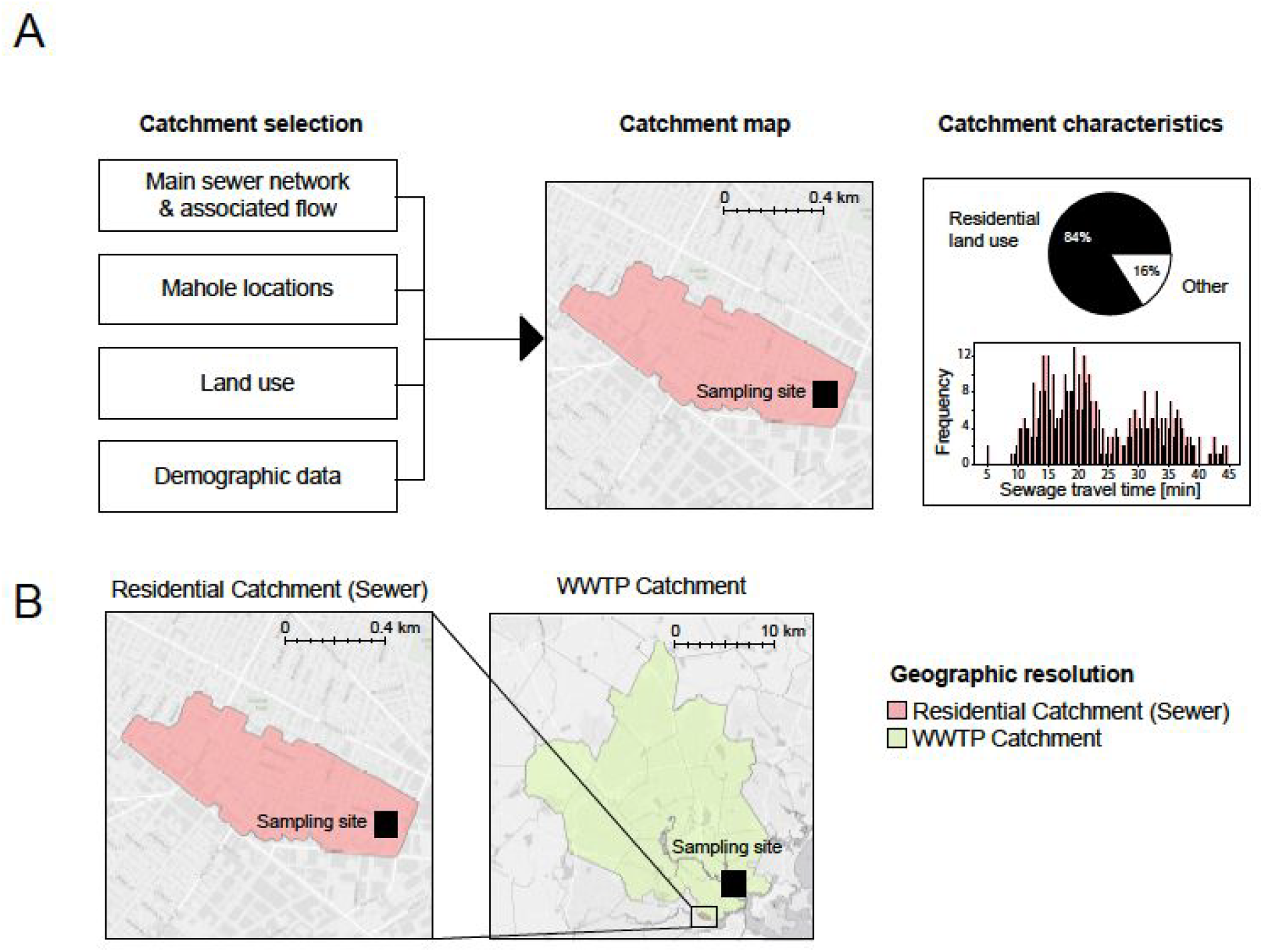
Overview of residential catchment and wastewater treatment plant (WWTP) sampling features. (A) Integrating sewer network, manhole location, land use, and demographic data allows for the selection of a sampling manhole that contains sewage from a residential catchment. Samples taken at this sampling location represent a catchment which is primarily residential and has a short travel time from the place of origin to the sampling location. (B) Comparison of the area of the residential catchment and WWTP catchment sampled in this study.

Metabolite relative abundances differed significantly between upstream and downstream sites. 34.5% of metabolites detected in both sewer and WWTP samples (n=245/710) had significantly different concentrations between these two sampling locations (q < 0.05; t-test with FDR correction). Of these differentially abundant metabolites, 73% (n=179/245) decreased or disappeared completely from wastewater collected at the WWTP, including many metabolites from human activity (see “Residential sewage microbiome and metabolome reflect human diurnal activity” section, Fig. 2A yellow dots, Fig. 3C yellow bars, Supp. Fig. 2).

**Figure 2.**
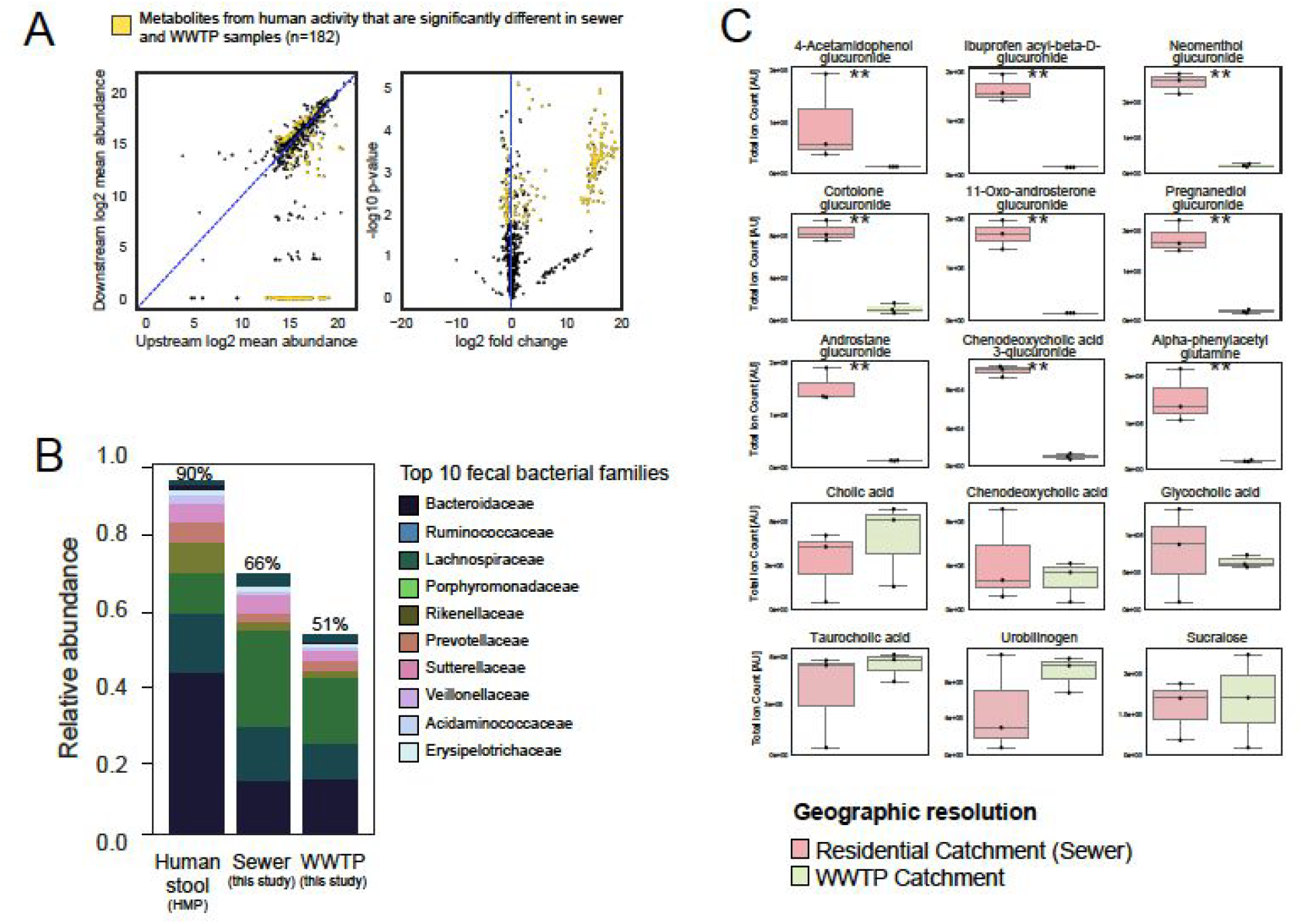
Comparison of metabolites and bacterial community composition between residential catchment (sewer) and WWTP catchment. (A) Left: log2 mean abundance of metabolites in upstream (sewer, n = 3) vs. downstream (WWTP, n = 3) samples. Right: log2 fold-change vs. log10 p-value for upstream vs. downstream comparison, log2 fold-change is for the ratio of upstream to downstream metabolites. Metabolites identified as relating to human activity from co-clustering analysis are indicated in yellow (see Fig 3C and Supp. Fig 5). Most of the significantly different human-associated metabolites between the two sites are detectable upstream but less or undetectable downstream. (B) The residential catchment (sewer) has a higher representation of the top 10 fecal bacterial families (i.e. families which represent 90% of fecal bacteria in the HMP, as in Newton et al. 2015) than the downstream site (WWTP). (C) Abundance of fifteen of the seventeen human activity-derived metabolites measured by targeted LC-MS/MS and identified at level 1 or 2 in upstream residential catchment (sewer, green) vs. downstream WWTP (red) (** p-value < 0.01, n = 3 samples per site, the remaining 2 metabolites are shown in Supp. Fig 4.). Human-associated compounds which are susceptible to biological and/or chemical degradation are readily present in the upstream sample but absent in WWTP samples (see top three rows of glucuronides), while more stable compounds show similar levels in both sampling locations (bottom two rows).

**Figure 3.**
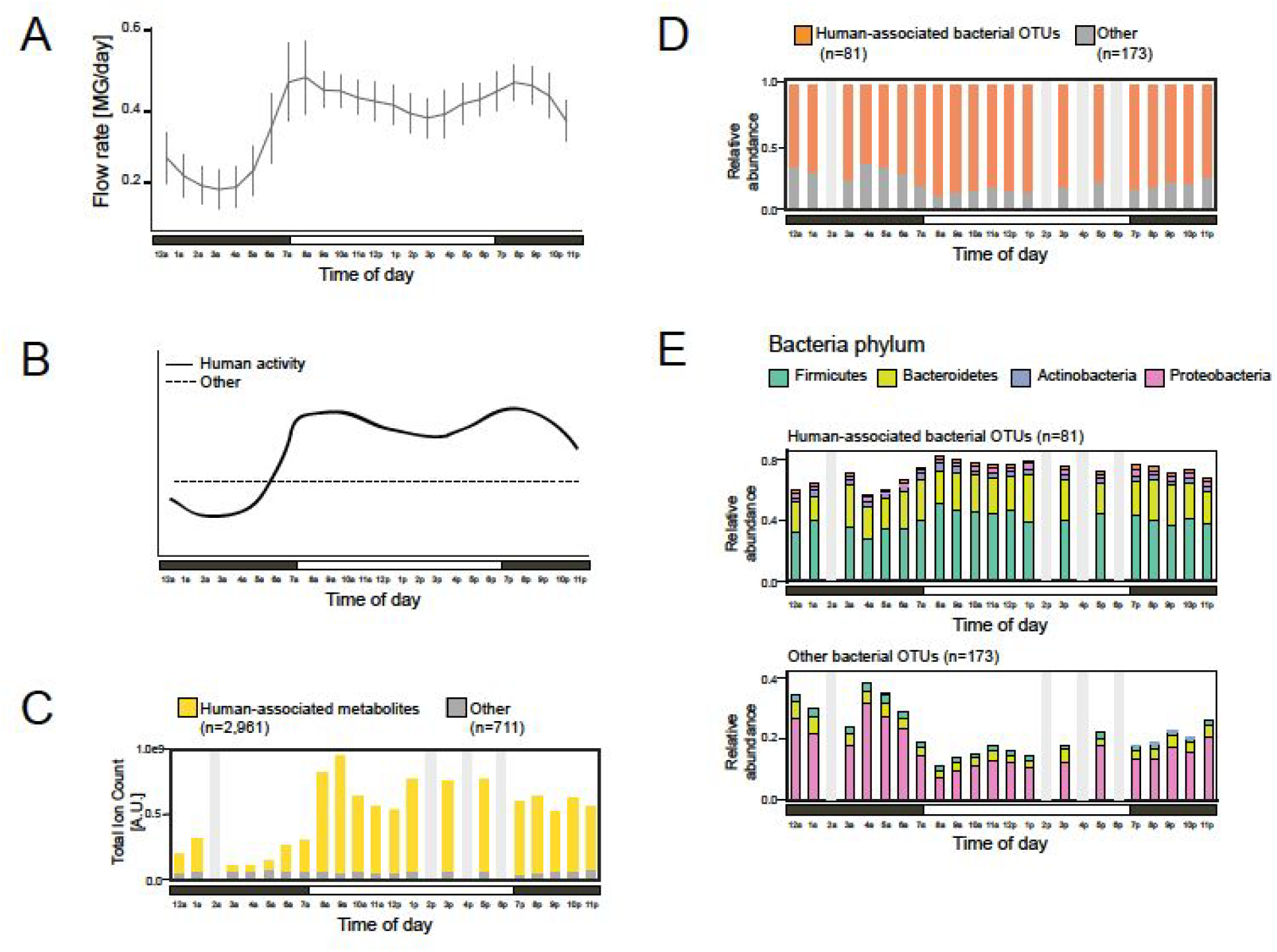
Dynamics of the metabolites and bacteria sampled in the residential catchment with hourly sampling for 24 hours. (A) Sewage flow rate varies across the day, with the lowest flow rate occurring at night when human activity is lowest. Flow is plotted as the mean and standard deviation of Tuesday-Thursday measurements taken every 15 minutes over four weeks. (B) Cartoon schematic showing the expected dynamics of human-derived (solid line) and other (dashed line) metabolites and bacteria given the flow rate dynamics and human diurnal activity. (C and D) Abundances of human-associated and non-human-associated metabolites (C) and OTUs (D) throughout the day. Human-associated groups were identified by clustering metabolite-OTU temporal correlations. Human-associated groups metabolites (C, yellow) and OTUs (D, orange) increase during the day and decrease at night, while non-human-associated metabolites and OTUs (gray) remain relatively constant throughout the 24-hour sampling period. 2AM, 2PM, 4PM, and 6PM samples did not meet QC standards and are not included (see Methods). (E) Phylum-level relative abundances over the course of the day for human-associated (top) and other (bottom) OTUs. The human-associated OTU cluster is dominated by human-derived bacteria (Firmicutes and Bacteroidetes), whereas the other cluster is dominated by environmental bacteria (Proteobacteria)

Biological signals also differed between upstream and downstream sites. Residential sewage contained a higher proportion of human-associated bacteria than WWTP sewage (Fig 2B). Human-associated bacteria were defined as the top ten bacterial families in human stool (as in Newton et al. 2015), representing 90% of the average stool from the Human Microbiome Project (Huttenhower 2012). On average, 66% of the microbial community from residential sewage was of human fecal origin by this definition, compared to 51% in the WWTP pump station samples (Fig. 2B), with reduced representation by the *Ruminococcaceae*, *Lachnospiraceae*, *Prevotellaceae*, *Veillonellaceae* and *Erysipelotrichaceae* families (Supp. Fig. 3). These results indicate that the abundance of human gut bacteria decreases in the sewer environment. We expect the rate of decrease to depend on sewer environmental conditions and to result in dramatically different amounts of human fecal bacteria in WWTPs. Consistent with this expectation, a previous study observed large variability in the relative amounts of human fecal bacteria in 71 WWTPs across the U.S. (1.59% – 69.2%; average 15%, Newton et al. 2015).

### Upstream sampling enables detection of human associated biomarkers *via* glucuronidation

Some metabolites are not typically detected in downstream analyses due to their reactivity in sewer systems. We hypothesized that such variants would be uniquely present in upstream sewage because they would be collected shortly after excretion. Glucuronides are chemical moieties added to exogenous molecules in the human liver to aid in their excretion and are thus specific indicators of human excretion (de Wildt 1999). Importantly, these charged moieties are often added to hydrophobic molecules that are not detectable by mass spectrometry. Thus, glucuronidation enables the quantification of molecules that would otherwise elude measurement. However, glucuronide moieties are labile and many bacteria express glucuronidase enzymes that hydrolyze them, reducing the signal at WWTPs (Pollet 2017, Jacox 2017). Thus, we hypothesized glucuronides present at upstream sites might be underrepresented or absent in WWTP samples.

We identified 21 glucuronidated compounds in our untargeted data, based on analytical standards as well as matches in mass/charge (m/z) and structural characteristics (MS2 spectra) (see Methods, Table 2). Glucuronides that were readily detectable in residential catchments were reduced or altogether absent in wastewater collected at the WWTP (nine examples shown in Fig. 2C, others shown in Supp. Fig. 4). By contrast, compounds that are resistant to biological and/or chemical degradation, such as the artificial sweetener sucralose, showed similar levels in samples from residential and WWTP catchments (six examples shown in Fig. 2C). Our data suggests that sampling upstream residential catchments can expand the diversity of molecules available for future public health surveillance applications (Daughton 2018, Jacox 2012).

**Table 2.** Metabolites identified in residential wastewater samples through untargeted metabolomics, and MS2 (level 2) or standard (level 1) matching. Identification levels are adapted from the Metabolomics Standards Initiative and described in the Methods.

| Compound | Identified metabolite | Retention time (min) | Measured m/z | Mono-isotopic mass | Description | ID Level |
| --- | --- | --- | --- | --- | --- | --- |
| Acetaminophen | p-Acetamidophenyl glucuronide | 3.32 | 326.088 | 327.095 | Pharmaceutical (pain killer) | 1 |
| alpha-N-phenylacetyl-L-glutamine (PAG) | alpha-N-phenylacetyl-L-glutamine (PAG) | 7.86 | 263.104 | 264.111 | Normal biofunction (waste nitrogen excreted in urine) | 1 |
| Chenodeoxycholic acid | Chenodeoxycholic acid | 20.55 | 391.286 | 392.293 | Bile acid | 1 |
| Cholic acid | Cholic acid | 17.63 | 407.281 | 408.288 | Bile acid | 1 |
| Fecal urobilinogen | Urobilinogen/Stercobilinogen | 12.3 | 589.303 | 590.31 | Normal biofunction (gut bacteria fermentation of bilirubin) | 1 |
| Glycocholic acid | Glycocholic acid | 16.03 | 464.302 | 465.309 | Bile acid | 1 |
| Sucralose | Sucralose | 8.42 | 430.984 | 396.015 | Diet (artificial sweetener) | 1 |
| Taurocholic acid | Taurocholic acid | 17.39 | 514.285 | 515.292 | Bile acid | 1 |
| Adrenaline | 3-Methoxy-4-hydroxyphenylglycol glucuronide | 6.65 | 359.098 | 360.106 | Normal biofunction (hormone and neurotransmitter) | 2 |
| Aldosterone | Tetrahydroaldosterone-3-glucuronide | 13.03 | 539.25 | 540.257 | Steroid hormone (mineralcorticoid) | 2 |
| Caprylic acid or Valproic acid | Octanoylglucuronide or Valproic acid glucuronide | 13.22 | 319.14 | 320.147 | Diet (milk) or pharmaceutical (epilepsy) | 2 |
| Cheno/deoxycholic acid | Cheno/deoxycholic acid 3-glucuronide | 16.05 | 567.318 | 568.325 | Bile acid | 2 |
| Cholic acid | Cholic acid glucuronide | 15.43 | 583.313 | 584.32 | Bile acid | 2 |
| Cholic acid | Cholic acid glucuronide | 13.57 | 583.313 | 584.32 | Bile acid | 2 |
| Cortolone | Cortolone-3-glucuronide | 13.08 | 541.266 | 542.273 | Steroid hormone | 2 |
| Dextrophan | Dextrophan glucuronide | 6.66 | 432.203 | 433.21 | Pharmaceutical (cough suppressant) | 2 |
| Ferulic acid | Ferulic acid 4-O-glucuronide | 8.89 | 369.083 | 370.09 | Diet (plants) | 2 |
| Ferulic acid | Dihydroferulic acid 4-O-glucuronide | 8.61 | 371.099 | 372.106 | Diet (plants) | 2 |
| Glycochenodeoxycholic acid | Glycochenodeoxycholic acid 3-glucuronide | 16.23 | 624.339 | 625.346 | Bile acid | 2 |
| Ibuprofen | Ibuprofen acyl glucuronide | 16.34 | 381.156 | 382.163 | Pharmaceutical (pain killer) | 2 |
| Menthol | Neomenthol glucuronide | 14.88 | 331.176 | 332.184 | Diet (plants) and personal care products | 2 |
| Mycophenolic acid | Mycophenolic acid glucuronide | 12.42 | 495.151 | 496.158 | Pharmaceutical (immunosuppressant) | 2 |
| Nicotine | trans-3-Hydroxycotinine glucuronide | 25.96 | 367.114 | 368.122 | Recreational drug | 2 |
| p-cresol | p-cresol glucuronide | 9.31 | 283.082 | 284.09 | Normal biofunction (gut bacteria fermentation) | 2 |
| Progesterone | Pregnanediol glucuronide | 17.36 | 495.297 | 496.304 | Steroid hormone (progestagen, sex hormone) | 2 |
| Testosterone | Testosterone glucuronide or Dehydro(epi/iso)androst erone 3-glucuronide | 15.48 | 463.234 | 464.241 | Steroid hormone (androgen, sex hormone) | 2 |
| Testosterone | Androstane glucuronide | 16.37 | 465.25 | 466.257 | Steroid hormone (androgen, sex hormone) | 2 |
| Testosterone | 11-Oxo-androsterone glucuronide | 14.03 | 479.229 | 480.236 | Steroid hormone (androgen, sex hormone) | 2 |
| Testosterone | 11-beta-Hydroxyandrost erone-3-glucuronide | 13.63 | 481.245 | 482.252 | Steroid hormone (androgen, sex hormone) | 2 |

**Figure 4.**
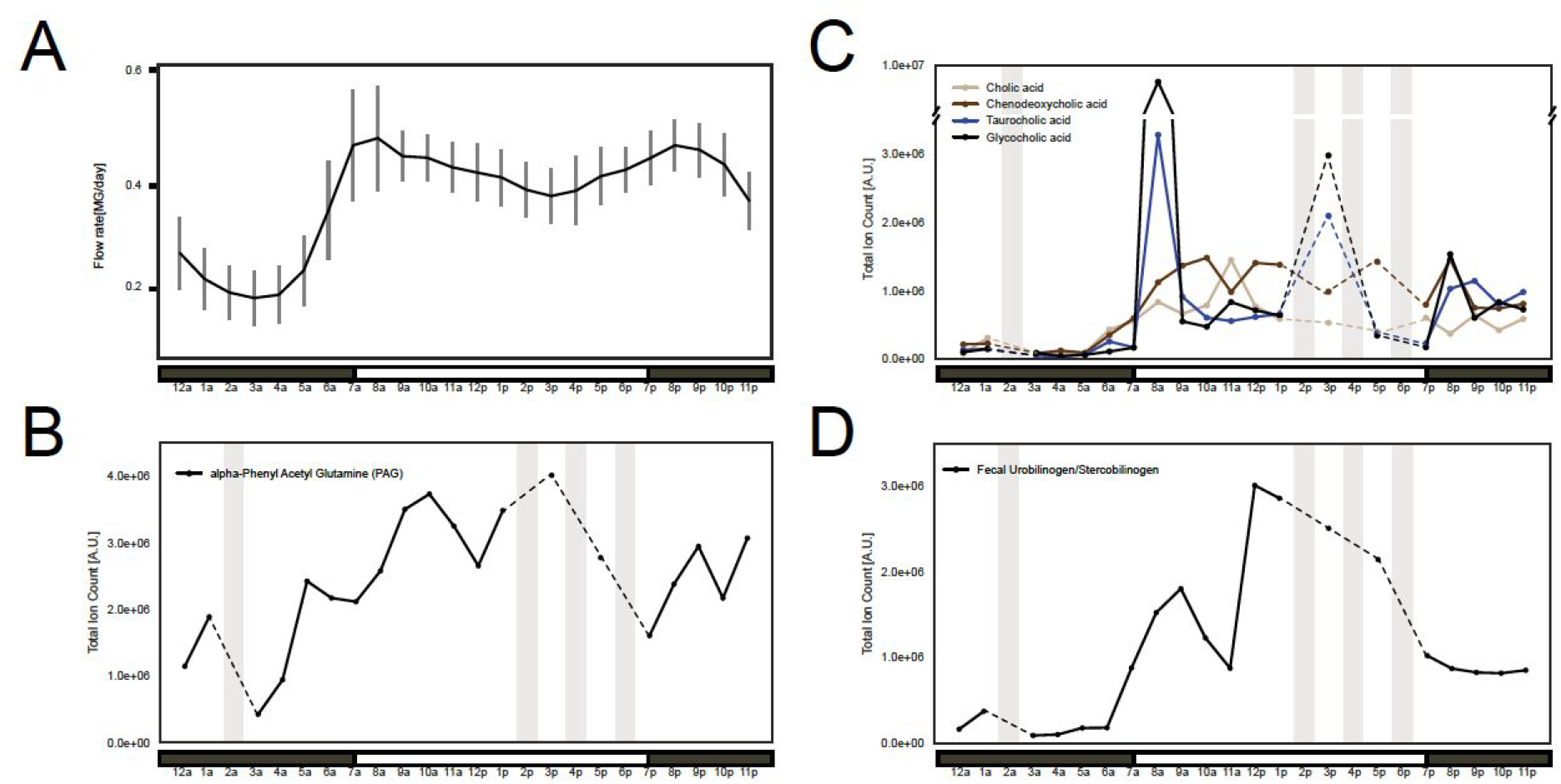
Daily dynamics of confirmed fecal and urinary metabolite markers. (A) Upstream residential catchment sewage flow rate (flow rate data is plotted as described in Fig 3A and Methods). (B) Abundance of urinary metabolite marker alpha-N-phenylacetyl-L-glutamine (PAG). (C) Abundance of four bile acids (conjugated: taurocholic & glycocholic, unconjugated: cholic and chenodeoxycholic). (D) Abundance of fecal metabolite marker urobilinogen/stercobilinogen. Dashed lines between solid lines indicate interpolated data. Metabolites shown were confirmed by analytical standard (level 1).

### Residential sewage microbiome and metabolome reflect human diurnal activity

Chemical and bacterial biomarkers detected in residential sewage reflect the diurnal cycles of human activity, as approximated by changes in average sewage flow rates collected over four weeks (Fig. 3A and B, Ort 2010). We hypothesized that human-associated metabolites and bacteria would increase during the day, when the contributing population would be more active, and decrease at night, whereas the environmental background would remain relatively constant (Fig. 3B). To test this hypothesis, we collected hourly samples over 24 hours from the residential catchment and produced 16S rDNA sequencing and untargeted metabolomics profiles. Metabolomics data from four time points did not pass quality control filters and these were excluded from our analysis (2AM, 2PM, 4PM, 6PM, see Methods).

Clustering metabolites and bacterial taxa enabled us to identify components associated with human activity. We calculated the Spearman correlations between metabolites (n=3,672) and bacterial taxa (n=254) and co-clustered metabolites and OTUs based on this correlation matrix. One co-cluster of metabolites (n=2,961/3,672) and bacteria (n=81/254) (dashed box, Supp. Fig. 5) had strong positive correlations to each other and increased during the day and became much less abundant at night. We identified these as putative “human-associated” features, and they included most of the metabolic features (mean abundance = 80%, yellow bars Fig. 3C) and bacteria in the sewer (mean relative abundance = 78%, orange bars Fig. 3D).

We next asked whether these metabolites and bacteria were indeed primarily human-associated by annotating the most abundant metabolites (see Methods). We found that the “human-associated” cluster was statistically enriched in human metabolites such as glucuronide compounds, hormones, bile acids and pharmaceuticals (Fisher’s exact test p-value = 1.4e-10, Table 3). Bacteria which clustered with the “human-associated” group of metabolites included 81 bacterial OTUs primarily from the *Firmicutes* and *Bacteroidetes* phyla, which are major components of the human gut flora (Fig. 3E, Huttenhower 2012). 65% of bacteria in the “human-associated” bacterial cluster belonged to the top ten most abundant bacterial families in human stool (Supp. Fig. 6).

**Table 3.** Metabolites in the human-associated cluster are enriched in metabolites derived from human activity (Fisher’s exact test, p-value = 1.4e-10). Subcategories of identified metabolites are indicated in italics; totals are bolded. Levels are adapted from the Metabolomics Standards Initiative (Sumner 2007) and described in the Methods.

| Type of identification | "Human-associated" cluster | "Other" cluster | Total |
| --- | --- | --- | --- |
| **Identified metabolites** | **308** | **24** | **332** |
| *Urine metabolites (Level 1)* | 2 | 0 |  |
| *Bile acids (Level 1)* | 4 | 0 |  |
| *Sucralose (Level 1)* | 1 | 0 |  |
| *Glucuronides (Level 1 or 2)* | 22 | 0 |  |
| *Human Metabolome DB (Level 3)* | 279 | 24 |  |
| **Unknown metabolites** | **2,653** | **687** | **3,340** |
| **Total** | **2,961** | **711** | **3,672** |

The cluster of non-“human-associated” metabolites (n=711) and bacterial OTUs (n=173), on the other hand, likely reflect bacteria and metabolites sourced from the environmental background rather than from human activity. These metabolites had few matches to human compounds and remained constant throughout the 24-hr sampling period, likely representing chemicals derived from the environment or sewer biogeochemistry. Similarly, this bacterial cluster was primarily composed of *Proteobacteria* (Fig. 3E), which are not major members of the human gut flora. Indeed, human fecal bacterial families comprised only 1.4% of the abundance of this cluster (Supp. Fig. 6). The relative abundance of the bacteria in this cluster increased at night, but this likely reflects the fact that there were fewer bacteria of fecal origin rather than an increase in environmental bacteria, since a similar increase in environmental metabolites was not observed (Fig. 3C and D, gray bars).

### Identification of urinary and fecal biomarkers

Urinary and fecal biomarkers supplement sewage flow rate as a more direct measure of human activity in sampling optimization, and can be used as a normalization factor for specific quantities of interest (Ort 2010). For example, drugs and pharmaceutical products are mainly excreted in urine, while certain infectious diseases like polio virus are shed in stool. To the extent that feces or urine is concentrated at certain times of day, sampling efforts could be matched to these hours to reduce the logistical burden of continuous sampling. In particular, we hypothesized that fecal matter might be concentrated in a reduced set of hours.

We identified and confirmed metabolites corresponding to human urinary and fecal markers through a combination of analytical standards and m/z plus MS2 matching, according to the protocols in the Metabolomics Standards Initiative (MSI, Sumner 2007). We prioritized mass spectral features with high abundance across all samples, and also in a few samples of interest (8 am and 1 pm samples, which contained more total metabolites and which we expected to correlate with human behavior). We screened the top m/z values with METLIN and MetFrag to characterize putative human-derived metabolites (Smith 2005, Wolf 2010). We compared all m/z values to the Human Metabolite Database (HMDB) database and used MetFrag to compare the observed MS2 fragmentation patterns of potential glucuronide compounds with their expected fragmentations (see Methods; Wishart 2012, Wolf 2010, Ashcroft 1995). When commercially available, we obtained analytical standards to confirm our identifications.

The most abundant feature in our dataset was identified (level 1 confidence, MSI, Sumner 2007) as alpha-N-phenylacetyl-L-glutamine (PAG), an end product of phenylalanine metabolism excreted in urine (Seakins 1971, Stein 1954). Previous studies on human physiology have found that PAG excretion remains constant in response to diet and does not exhibit diurnal cycling in individual patients (Seakins 1971). Thus, the abundance of PAG in our sample likely correlates with the total volume of urine in sewage at a given time. Urobilinogen, a hemoglobin breakdown product excreted primarily in stool (Balikov 1957), was also identified (level 1 confidence). We identified four bile acids (chenodeoxycholic acid, cholic acid, glycocholic acid, taurocholic acid; level 1 confidence). We considered these compounds as putative fecal markers, as they represent primary conjugated (produced in the liver) and unconjugated bile acids (acted on by bacterial bile salt hydrolases) that aid in digestion (Hofmann 1989).

Urinary and fecal metabolites tracked general diurnal human activity, but their dynamics differed from each other (Fig. 4). Both types of metabolites decreased at night, with the fecal markers almost completely disappearing (Fig 4C and D) while the urinary marker remained detectable (Fig 4B), albeit at reduced levels. Fecal markers also showed large deviations in the 8 am and 3 pm samples, and were slightly elevated throughout the evening (8 – 10 pm, Fig 4C and D). In contrast, the urinary markers increased in the morning and stayed relatively consistent throughout the day (Fig 4B). These patterns correspond with known patterns of human behavior: during the day, urination remains relatively constant across the entire population, while defecation spikes in the morning right after waking and in the early afternoon (Heaton 1992). Thus, sampling continuously throughout the day may be needed to capture the urinary fraction (see PAG, Fig. 4B), while fecal surveillance could be targeted to early mornings (see bile acids, Fig. 4C).

### Sewage reveals dynamics of bile acid production in humans

Human production and excretion of bile acids is a complex physiological process, with conjugated and unconjugated bile acids exhibiting related but different responses to meals and to specific types of dietary intake (Hofmann 1989). The daily dynamics of fecal bile acid excretion is not well studied in humans, in part because it is difficult to access temporally-resolved data: humans defecate only 1-3 times per day on average or irregularly (Heaton 1992). Our 24-hour samples provide a unique opportunity to study the mix of bile acids on a population level.

We previously reported day-to-day variability in the types and quantities of bile acids produced in a single individual (Kearney 2018). Using our sewage aggregate data, we found additional variability in the types of bile acids excreted at different times of day. Levels of unconjugated cholic and chenodeoxycholic acids remained relatively constant during the day but conjugated cholic acids (glycocholic and taurocholic acid) showed more variation, with a notable spike in the morning (Fig 4C). These spikes in conjugated cholic acids may reflect an aggregate population-level physiological response to meals, or simply the sampling variability of residential sewage. Estimates of per sample population size are needed to discriminate these cases. In either case, these data suggest that analyzing physically aggregated biological data could provide an alternative avenue by which to study human physiology.

### Residential sewage contains potential indicators of community health

Novel potential biomarkers of population health and behavior in our metabolomics data suggest that residential sewage is a promising source of community health indicators (Daughton 2018, Metzler 2008). 332 out of 3,672 untargeted metabolomic features were putatively identified in the HMDB database. Within the putatively-annotated glucuronides, we found human physiology biomarkers, compounds derived from drugs and pharmaceuticals, and dietary metabolites (Fig. 5 and Table 2). Physiological biomarkers included glucuronidated hormones and steroids such as adrenaline, aldosterone, and cortolone (Table 2), and other endogenous markers of key physiological processes could likely be identified (Daughton 2018). We found many examples of glucuronidated drug compounds, including over-the-counter painkillers (acetaminophen and ibuprofen glucuronide), immunosuppressants (mycophenolic acid glucuronide), and recreational drugs (3-hydroxycotinine glucuronide, a metabolite of nicotine). Such compounds could be measured to track human consumption, identify communities with high use of pharmaceuticals, or evaluate the effectiveness of drug take-back programs (Egan 2017). Finally, some metabolites mapped to dietary markers like sucralose, an artificial sweetener, and metabolites of ferulic acid, a phenolic acid abundant in fruits, vegetables, and beverages like juice and coffee (Piazzon 2012).

**Figure 5.**
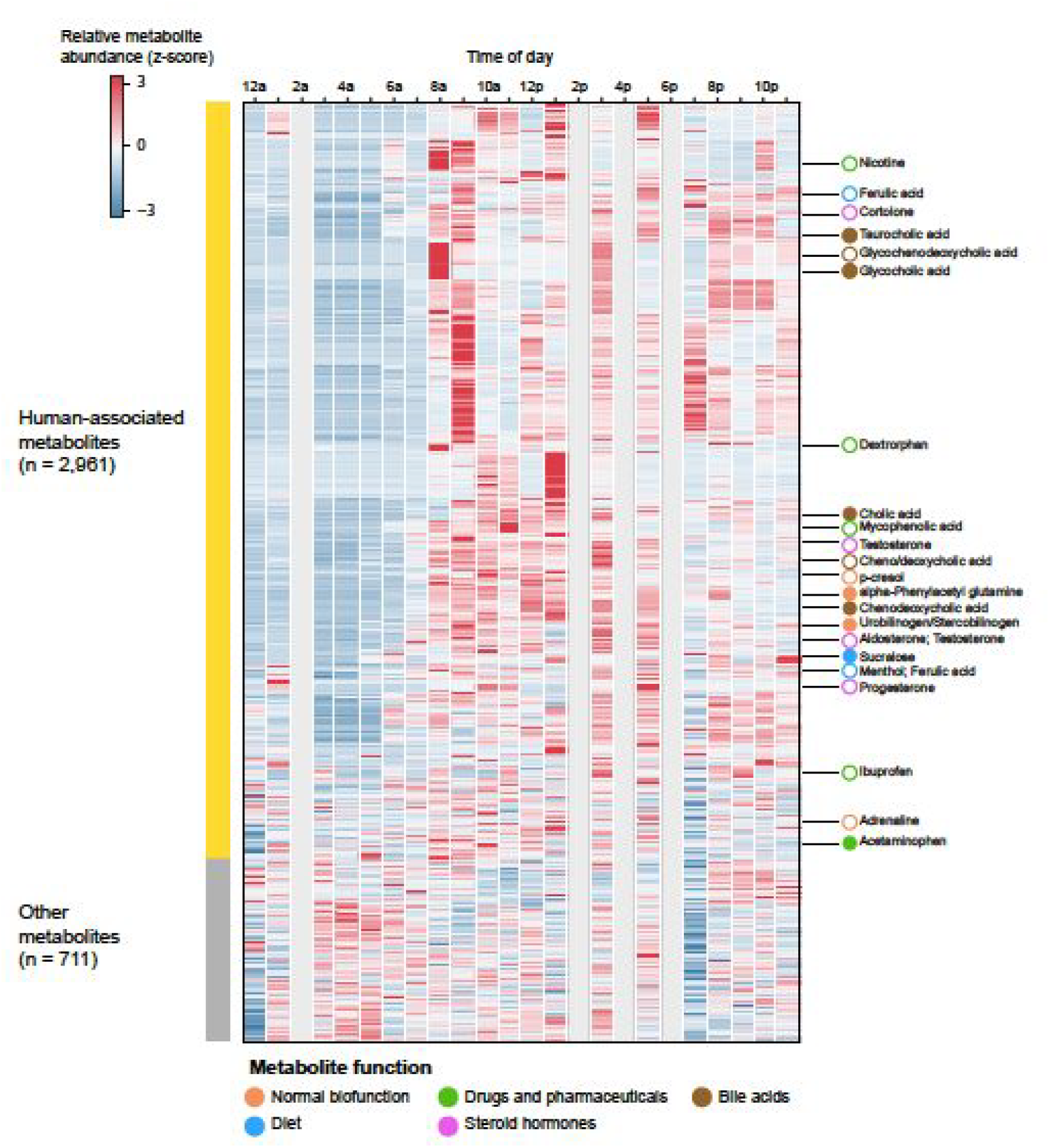
Normalized metabolite abundance over time. Each metabolite feature (rows) is Z-scored across time (columns). Metabolites are ordered according to the co-clustering between metabolites and OTUs, as described in the Methods. Metabolites in the human-associated groups are shown in yellow, others are in gray (left bar). Putatively identified metabolites are indicated to the right of the heatmap, with filled-in circles indicating level 1 or 2 identifications (confirmed by analytical standard and/or MS2) and empty circles indicating level 3 identifications (mz match only). All indicated metabolites (except acetaminophen) fall into the human-associated cluster.

## Discussion

Here, we present a framework to adapt the practices of wastewater-based epidemiology to the unique needs of city-level public health officials. We show that upstream sampling and leveraging human diurnal activities is key to unlocking much of the potential of sewage surveillance. Untargeted genomic and metabolomic data, rather than a targeted approach to measure a few known biomarkers, allowed us to uncover a wide range of biomarkers that we would not have anticipated *a priori*.

Such biomarkers may reflect a wide range of human activities and could also be used for a wide range of policy applications. For example, analyzing the amounts of un-glucuronidated prescription opioids or antibiotics could be used to evaluate the effectiveness of drug take-back programs. Bacterial or chemical biomarkers of nutrition intake and diet could be used to study food deserts and the impact of programs to increase the availability of fresh foods in certain neighborhoods. Marijuana metabolites and plant strains could be measured in sewage to measure the impact of cannabis legalization roll-outs across different counties or states. Residential sewage monitoring could also alert city officials of infectious disease outbreaks before a corresponding uptick in clinical cases, enabling earlier intervention in the affected communities. Finally, mining residential sewage can also uncover behaviors not effectively measured by other means. For example, identifying communities with high rates of unreported non-fatal opioid overdoses could be possible by finding catchments with high levels of naloxone-associated metabolites but low reported overdose rates (Walley 2013, Beeson 2018).

Despite the exciting results presented here, we recognize key limitations and areas for future work. First, limited grab samples show strong variability (Supp. Fig 2, Lai 2011), suggesting the need for development of continuous sampling methods, ideally proportional to flow and possibly urine/feces content (Ort 2010). In addition, estimates of per sample population size could allow more quantitative inferences, such as estimating not only presence of an emerging pathogen, but its incidence within a given population (Daughton 2012, O’Brien 2013, van Nuijs 2011). In addition, identifying molecules was a significant bottleneck in our analysis, thus larger collections of chemical standards and more sophisticated computational approaches are needed to expand the set of quantifiable metabolites. As new tools and methods are developed, we expect that the data generated in this and future studies could be re-analyzed and mined for additional novel insights. To enable such re-analyses, our data is available in public databases (see Methods).

Finally, data privacy and ethics should be considered in all stages of development and deployment. Sewage is a physical aggregate of multiple individuals, and so does not share many of the same re-identification concerns as human genomics and microbiome studies (Hall 2012, Prichard 2014, Lowrance 2007, Franzosa 2015). However, concerns about disparate impact and data misuse are real and should be addressed with continual input and feedback from all stakeholders, including not just the scientists and engineers developing these platforms for deployment but also community leaders, sociologists, and policy and privacy experts.

The last decade has seen exponential growth of a "data economy", which has transformed almost every aspect of our lives. Yet this revolution has focused almost entirely on digital data, while very little has been done to capture biological and chemical data. Upstream sewage analysis represents a unique opportunity to tap into this wealth of data for public health and policy evaluation impact, ensuring that public health practitioners and other public officials also benefit from the increased interest in and development of “big data”, ‘omics, and precision medicine tools and technologies.

## Materials and Methods

### Selection of Residential Catchment sampling site

The sampling site was identified by cross-referencing Cambridge’s wastewater network maps, which contained the layout of pipes and manholes, as well as flow direction, with demographic and land use data. We selected a location where the upstream catchment fulfilled three major requirements: 1) a land use of over 80% residential so that we avoid transient populations and can characterize the demographics of the catchment population accurately, 2) a total catchment population >5,000 people to provide an anonymous reading of the community, and 3) the wastewater travel time from the furthest point in the catchment is <60 minutes to preserve the integrity of the sample. Average sewage flow rate at the selected manhole was calculated by measuring sewage flow rate every 15 minutes with a commercial flow meter over four weeks, then averaging mid-week measurements only (Tuesday-Thursday). Average sewage flow rate at the WWTP pump station was provided by their engineering staff.

### Wastewater collection and processing

For the time series analysis, 24 time-proportional samples were collected. Grab wastewater samples (500 mL each) were taken every hour from 10 AM on Wednesday April 8 2015 to 9 AM on Thursday April 9 2015. There was light rain during most of the day (on average <0.01 in) with peak precipitation between 11pm and 12am (0.09 in). For the comparison between the residential and the WWTP catchments, three grab samples (500 mL each) were collected from both the residential manhole and a pump station directly upstream of the WWTP on Thursday October 15 2015, and approximately at the same time (12pm – 2pm). There was no rain in the geographical area represented by the residential and WWTP catchments on this day. Sewage at the manhole was collected with a commercial peristaltic pump (Global Water) sampling at a rate of 100 mL/min. Sewage at the WWTP pump station was collected with a commercial sampling pole.

For all samples, 200 ml of collected sewage were filtered through 10-μm PTFE membrane filters. 150 ml of the outflow were filtered through 0.2-μm PTFE membrane filters. 100 ml of the final outflow were acidified with concentrated HCl (Optima grade, Fisher Scientific) to pH 3.0 and frozen at −80 degrees Celsius until metabolomics analysis. PTFE membrane filters were kept in RNAlater at −80 degrees Celsius until DNA extraction. Deionized water was processed through the same filtration system to generate blank filtrates and blank PTFE membranes. Four blanks were generated in total, one was collected at the start before any wastewater had passed through the system, another one at the end, and two more were spread over the processing time. The lab filtration system consisted of a Masterflex peristaltic pump (Pall), Masterflex PharMed BPT Tubing (Cole-Palmer), 47 mm PFA filter holders (Cole-Palmer) and 47mm PTFE Omnipore filter membranes (Millipore). Both the tubing and filter holders were previously cleaned with HCl (10% v/v) and ultrapure deionized water. The tubing and filter holders were flushed with 10% bleach and with distilled water, for 10 minutes each after every wastewater sample was processed.

### DNA extraction and amplicon based Illumina sequencing of 16S rRNA genes

0.2-μm filter membranes were thawed in ice. RNAlater was removed and filters were washed with phosphate-buffered saline (PBS) buffer twice. DNA was extracted from each filter with Power Water extraction kit (MO BIO Laboratories Inc.) according to manufacturer’s instructions. The only modification was that DNA from up to three filters from the same sample were pooled into the same DNA-binding column. Paired-end Illumina sequencing libraries were constructed using a two-step PCR approach targeting 16S rRNA genes previously described (Preheim 2013). All paired-end libraries were multiplexed into one lane and sequenced with paired end 300 bases on each end on the Illumina MiSeq platform at the MIT Biomicro Center.

### Processing of 16S rRNA gene sequencing data

Raw data was processed with an in-house 16S processing pipeline (https://github.com/thomasgurry/amplicon_sequencing_pipeline). The specific processing parameters can be found in Supplementary Table 1. To assign OTUs, we clustered OTUs at 99% similarity using USEARCH (Edgar 2010) and assigned taxonomy to the resulting OTUs with the RDP classifier (Wang 2007) and a confidence cutoff of 0.5. For each dataset, we removed samples with fewer than 100 reads and OTUs with fewer than 10 reads. We calculated the relative abundance of each OTU by dividing its value by the total reads per sample. We then collapsed OTUs by their RDP assignment up to genus level by summing their respective relative abundances. OTUs without annotation at the genus-level were combined into one “unknown” genus.

### Untargeted metabolomics analytical methods

Acetonitrile and deuterated biotin (d2-biotin) were added to the 0.2 um-filtrate (acetonitrile final concentration was 5%, deuterated biotin was 0.05 µg ml^−1^). The resulting solution was analyzed using liquid chromatography (LC) coupled via electrospray ionization (negative ion mode) to a linear ion trap-7T Fourier-transform ion cyclotron resonance hybrid mass spectrometer (Thermo Scientific, FT-ICR MS; LTQ FT Ultra). Chromatographic separation was performed using a Synergi Fusion C18 reversed phase column (2.1 × 150mm, 4um, Phenomenex). Samples were eluted at 0.25 mL/min with the following gradient: an initial hold at 95% A (0.1% formic acid in water): 5% B (0.1% formic acid in acetonitrile) for 2 minutes, ramp to 65% B from 2 to 20 minutes, ramp to 100% B from 20 to 25 min, and hold at 100% B until 32.5 minutes. The column was re-equilibrated for 7 min between samples at 95% solvent A. The injected sample volume was 20 uL. Full MS data were collected in the FT-ICR cell from m/z 100-1000 at 100,000 resolving power (defined at 400 m/z). In parallel to the FT acquisition, MS/MS (MS2) scans of the four most intense ions were collected at nominal mass resolution in the linear ion trap (LTQ). Samples were analyzed in random order with a pooled sampled run every six samples in order to assess instrument variability.

### Untargeted metabolomics raw data processing

Raw XCalibur files were converted to mzML files with MSConvert (threshold = 1000 for the negative ion mode files) (Chambers 2012). Peaks were picked using the function xcmsSet from the R package xcms, with the following parameters (negative mode): method = ‘centWave’, ppm = 2, snthresh = 10, prefilter = (5, 1000), mzCenterFun = ‘wMean’, integrate = 2, peakwidth = (20, 60), noise = 1000, and mzdiff = −0.005 (Smith 2006). To align retention times, we used the retcor.obiwarp function with the following parameters: plottype=‘deviation’, profStep=0.1, distFunc=‘cor’, gapInit=0.3, and gapExtend=0.4. To group peaks from different samples we used the group.density function with the following parameters: minfrac=0, minsamp=1, bw=30, and mzwid=0.001. To integrate areas of missing peaks, we used the fillPeaks function with method=‘chrom’.

Sample-wise QC was performed based on the spiked in d2-biotin (features identified as Cl adduct: mz = 281.070 and rt = 8.6 min; and H adduct: mz = 245.077 and rt = 8.5 min). We discarded samples with lower recovery of d2-biotin, as we considered this evidence of ion suppression and thus reduced confidence in the peak area values for the remaining features. Specifically, the 2 am, 2 pm, 4 pm, and 6 pm had levels of d2-biotin more than one standard deviation below the mean of all samples, and were discarded from further analyses.

We also performed feature-wise QC on the metabolomics data. For the 24-hour sampling data, we discarded peaks which were present in any of the 4 blank samples or which were absent in any 7 of the pooled samples. We determined each peak’s coefficient of variation (CV) within the pooled samples by calculating its standard deviation divided by the mean, and discarded peaks with a CV greater than 0.35. We also discarded any peaks which were present in fewer than 4 of the time samples. For the upstream/downstream samples, we discarded any features present in either of the 2 blanks.

### Metabolite assignments

#### Mass matching (level 3 confidence)

We assigned putative metabolites according to the confidence levels proposed by the Metabolomics Standards Initiative (Sumner 2007). A level-3 confidence putative classification describes molecules whose observed masses match calculated exact mass values. Here we compared 397 of the most abundant mass features with METLIN (Smith 2005) and putatively characterized 126 (ppm error <= 2). We also programmatically mapped all metabolite m/z values to the Human Metabolome Database (HMDB) database (Wishart 2013, https://github.com/cduvallet/match_hmdb, database as of May 2016). Untargeted m/z values were first converted to their expected neutral mass (i.e. [M-H]- or [M+Cl]-ions), and then scanned against the neutral masses of all HMDB compounds, with an error tolerance of 1 ppm. All HMDB compounds within the error tolerance were returned. Results are shown in Supplementary Table 2. An m/z with a single hit in HMDB (“hit_type” column = hmdb_name) shows the chemical name of the HMDB entry and its HMDB ID. When an m/z had multiple HMDB hits, the feature was named with the common chemical taxonomic classification (“hit_type” column).

#### Matches by fragmentation spectra (level 2 confidence)

For a level-2 putative annotation, we compared observed fragmentation spectra (MS/MS) with *in silico* or database spectra. Here we focused on putative glucuronide derivatives from our level 3 comparisons (from m/z hits through METLIN and HMDB searching). MS/MS spectra were extracted from individual mzML files via MZMine (Pluskal 2010) and pymzML (Bald and Barth 2012) and matched to predicted *in silico* fragmentation patterns from MetFrag (Wolf 2010). We considered a glucuronide to be putatively annotated if the expected glucuronide was among the top MetFrag hits and the observed fragments included peaks corresponding to the un-glucuronidated compound, diagnostic glucuronide derivatives (m/z = 113, 157, 175), or other diagnostic peaks (parent compound minus a CO2 and/or OH) (Ashcroft 1995).

#### Identifications (level 1 confidence)

For a level 1 identification, putative annotations or characterizations must be confirmed through matches of m/z, retention time and fragmentation spectra of commercial standards analyzed on our platform (Sumner 2007). For these efforts, we purchased several metabolites (Table 2, ID level 1). Each standard was analyzed individually in pure solvent as well as spiked into the 1 pm and 2 am samples. We extracted the mz and retention times corresponding to our standards from the aligned feature table (level 3 m/z match as described above). We found the corresponding peaks in the standard-only runs via pymzML (m/z error between feature and peak in standard-only run = 0.005, RT error between feature and peak = 30 seconds) and also confirmed them by identifying their corresponding m/z and retention time in the spike-in samples. Next, we extracted the MS/MS spectra for each standard from its standard-only run and compared it to a corresponding MS/MS in an unspiked sample. For urobilinogen, the standard-only run did not trigger an MS/MS, so we compared its MS/MS from the spiked sample with one from a non-spiked sample. These analyses were performed through a combination of MZMine (Pluskal 2010) and pymzML (Bald and Barth 2012). Detailed results from the level 1 identifications are available in Supplementary File 1.

Metabolites in the upstream/downstream and 24-hour sampling experiment runs were matched based on their mz and retention time (ppm error < 5 for mz, ΔRT < 30 seconds). 867 metabolites were found to match between the two datasets. Of the 29 molecules identified via MS2 or standard matching (level 1 or 2) in the 24-hour dataset, 17 were also found in the upstream/downstream dataset.

## Code and data availability

Code to reproduce these figures are available at https://github.com/mmatus/24hr_publication. Raw and processed microbiome and metabolomics datasets will be made available upon publication in the respective databases (SRA, MetaboLights, and Zenodo).

## Acknowledgements

We thank The Cambridge Department of Public Works, in particular Jim Wilcox from the City of Cambridge, and Sam Lipson from the Cambridge Public Health Department for all their support in selecting, accessing and manning manholes. We are indebted to all researchers from the Alm Lab, Senseable City Lab, and Runstadler Lab who volunteered their time to help us collect and process sewage non-stop through one day, and for Herbie from Cambridge Public Works for providing his good humor and professional insights during our many sampling trips. We thank the generous sponsorship of the Kuwait Foundation for Advancement of Sciences, and the MIT Center for Microbiome Informatics and Therapeutics provided support for computational resources. We thank the WHOI FT-MS facility for sample analysis and Krista Longnecker for training in metabolomics data analysis and discussions throughout the study.

## References

Arcaya, M., Reardon, T., Vogel, J., Andrews, B. K., Li, W., & Land, T. (2014). Peer reviewed: Tailoring community-based wellness initiatives with latent class analysis—Massachusetts Community Transformation Grant Projects. Preventing chronic disease, 11.

Arnaud, Celia H. To monitor the health of cities’ residents, look no further than their sewers. (2018). Chemical & Engineering News, 96(18).

Ashcroft, A. E., Major, H. J., Lowes, S., & Wilson, I. D. (1995). Identification of non-steroidal anti-inflammatory drugs and their metabolites in solid phase extracts of human urine using capillary electrophoresis–mass spectrometry. In Analytical Proceedings including Analytical Communications (Vol. 32, No. 11, pp. 459–462). Royal Society of Chemistry.

Australian Criminal Intelligence Commission. National Wastewater Drug Monitoring Program—Report 5, August 2018. (2018) https://www.acic.gov.au/publications/intelligence-products/national-wastewater-drug-monitoring-program-report. Accessed December 2018.

Bald, T., Barth, J., Niehues, A., Specht, M., Hippler, M. and Fufezan, C., 2012. pymzML-Python module for high-throughput bioinformatics on mass spectrometry data. Bioinformatics, 28 (7), pp.1052–1053.

Balikov, B., 1957. Urobilinogen Excretion in Normal Adults: Results of Assays with Notes on Methodology. Clinical chemistry, 3(3), pp.145–153.

Baz-Lomba, J.A., Salvatore, S., Gracia-Lor, E., Bade, R., Castiglioni, S., Castrignanò, E., Causanilles, A., Hernandez, F., Kasprzyk-Hordern, B., Kinyua, J. and McCall, A.K., (2016). Comparison of pharmaceutical, illicit drug, alcohol, nicotine and caffeine levels in wastewater with sale, seizure and consumption data for 8 European cities. BMC public health, 16(1), p.1035.

Beeson, J. (2018). Notes From the Field-ODMAP: A Digital Tool to Track and Analyze Overdoses. https://www.ncjrs.gov/App/Publications/abstract.aspx?ID=273927

Berchenko, Y., Manor, Y., Freedman, L.S., Kaliner, E., Grotto, I., Mendelson, E. and Huppert, A., (2017). Estimation of polio infection prevalence from environmental surveillance data. Science translational medicine, 9(383), p.eaaf6786.

Burgard, D., Banta-Green, C., and Field, J.A. 2014. Working Upstream: How Far Can You Go with Sewage-Based Drug Epidemiology? Environmental Science and Technology. 48 (3), pp 1362–1368.

Castiglioni, S., Zuccato, E., Crisci, E., Chiabrando, C., Fanelli, R. and Bagnati, R., 2006. Identification and measurement of illicit drugs and their metabolites in urban wastewater by liquid chromatography− tandem mass spectrometry. Analytical Chemistry, 78(24), pp.8421–8429.

Chambers, M. C., Maclean, B., Burke, R., Amodei, D., Ruderman, D. L., Neumann, S., … & Hoff, K. (2012). A cross-platform toolkit for mass spectrometry and proteomics. Nature biotechnology, 30(10), 918.

Daughton, C. G. (2012). Real-time estimation of small-area populations with human biomarkers in sewage. Science of the Total Environment, 414, 6–21.

Daughton, C.G., (2018). Monitoring wastewater for assessing community health: Sewage Chemical-Information Mining (SCIM). Science of The Total Environment, 619, pp.748–764.

de Wildt, S. N., Kearns, G. L., Leeder, J. S., & van den Anker, J. N. (1999). Glucuronidation in humans. Clinical pharmacokinetics, 36(6), 439–452.

Dunn, W. B., Broadhurst, D., Begley, P., Zelena, E., Francis-McIntyre, S., Anderson, N., … & Nicholls, A. W. (2011). Procedures for large-scale metabolic profiling of serum and plasma using gas chromatography and liquid chromatography coupled to mass spectrometry. Nature protocols, 6(7), 1060.

Edgar, R. C. (2010). Search and clustering orders of magnitude faster than BLAST. Bioinformatics, 26(19), 2460–2461.

Egan, K. L., Gregory, E., Sparks, M., & Wolfson, M. (2017). From dispensed to disposed: evaluating the effectiveness of disposal programs through a comparison with prescription drug monitoring program data. The American journal of drug and alcohol abuse, 43(1), 69–77.

Escobar, M. A. C., Veerman, J. L., Tollman, S. M., Bertram, M. Y., & Hofman, K. J. (2013). Evidence that a tax on sugar sweetened beverages reduces the obesity rate: a meta-analysis. BMC public health, 13(1), 1072.

Fahrenfeld, N. and Bisceglia, K.J. (2016) Emerging investigators series: sewer surveillance for monitoring antibiotic use and prevalence of antibiotic resistance: urban sewer epidemiology. Environmental Science Water Research & Technology, 2, 788–799.

Falbe, J., Thompson, H. R., Becker, C. M., Rojas, N., McCulloch, C. E., & Madsen, K. A. (2016). Impact of the Berkeley excise tax on sugar-sweetened beverage consumption. American journal of public health, 106(10), 1865–1871.

Franzosa, E.A., Huang, K., Meadow, J.F., Gevers, D., Lemon, K.P., Bohannan, B.J. and Huttenhower, C., 2015. Identifying personal microbiomes using metagenomic codes. Proceedings of the National Academy of Sciences, p.201423854.

Hall, W., Prichard, J., Kirkbride, P., Bruno, R., Thai, P. K., Gartner, C., … & Mueller, J. F. (2012). An analysis of ethical issues in using wastewater analysis to monitor illicit drug use. Addiction, 107(10), 1767–1773.

Hanna-attisha M, Lachance J, Sadler RC, Schnepp AC (2016) Elevated Blood Lead Levels in Children Associated With the Flint Drinking Water Crisis: A Spatial Analysis of Risk and Public Health Response. 106(2):283–290.

Heaton, K. W., Radvan, J., Cripps, H., Mountford, R. A., Braddon, F. E., & Hughes, A. O. (1992). Defecation frequency and timing, and stool form in the general population: a prospective study. Gut, 33(6), 818–824.

Hellmér, M., Paxéus, N., Magnius, L., Enache, L., Arnholm, B., Johansson, A., Bergström, T. and Norder, H., (2014). Detection of pathogenic viruses in sewage gave early warning on hepatitis A and norovirus outbreaks. Applied and environmental microbiology, pp.AEM-01981.

Hofmann, A.F., 1989. Enterohepatic circulation of bile acids. Handbook of Physiology. The Gastrointestinal System, 4, pp.567–596.

Huttenhower, C., Gevers, D., Knight, R., Abubucker, S., Badger, J. H., Chinwalla, A. T., … & Giglio, M. G. (2012). Structure, function and diversity of the healthy human microbiome. Nature, 486(7402), 207.

Initiatives for Addressing Antimicrobial Resistance in the Environment: Current Situation and Challenges. (2018). https://wellcome.ac.uk/sites/default/files/antimicrobial-resistance-environment-report.pdf

Jacox, A., Wetzel, J., Cheng, S.Y. and Concheiro, M., (2017). Quantitative analysis of opioids and cannabinoids in wastewater samples. Forensic Sciences Research, 2(1), pp.18–25.

Kaliner, E., Kopel, E., Anis, E., Mendelson, E., Moran-Gilad, J., Shulman, L.M., Singer, S.R., Manor, Y., Somekh, E., Rishpon, S. and Leventhal, A., (2015). The Israeli public health response to wild poliovirus importation. The Lancet Infectious Diseases, 15(10), pp.1236–1242.

Kearney, S.M., Gibbons, S.M., Poyet, M., Gurry, T., Bullock, K., Allegretti, J.R., Clish, C.B. and Alm, E.J., 2018. Endospores and other lysis-resistant bacteria comprise a widely shared core community within the human microbiota. The ISME journal, 12(10), p.2403.

Keshaviah, A., Gitlin, R., Cattell, L., Reeves, W., de Vallance, J., and Thornton, C. (2016). The Potential of Wastewater Testing for Rapid Assessment of Opioid Abuse (Research Brief). Princeton, NJ: Mathematica Policy Research. https://www.mathematica-mpr.com/our-publications-and-findings/publications/the-potential-of-wastewater-testing-for-rapid-assessment-of-opioid-abuse-research-brief

Keshaviah, A. (ed.). (2017). The Potential of Wastewater Testing for Public Health and Safety. Washington, DC: Mathematica Policy Research. https://www.mathematica-mpr.com/our-publications-and-findings/publications/the-potential-of-wastewater-testing-for-public-health-and-safety-special-report

Lai, F. Y., Ort, C., Gartner, C., Carter, S., Prichard, J., Kirkbride, P., … & Mueller, J. F. (2011). Refining the estimation of illicit drug consumptions from wastewater analysis: co-analysis of prescription pharmaceuticals and uncertainty assessment. Water research, 45(15), 4437–4448.

Lowrance, W. W., & Collins, F. S. (2007). Identifiability in genomic research. Science, 317(5838), 600–602.

Manor Y, Shulman LM, Kaliner E, Hindiyeh M, Ram D, Soer D, Moran-Gilad J, Lev B, Grotto I, Gamzu R, Mendelson E., (2014). Intensified environmental surveillance supporting the response to wild poliovirus type 1 silent circulation in Israel, 2013. Euro Surveillance.19(7), p.20708

Metzler, M., Kanarek, N., Highsmith, K., Straw, R., Bialek, R., Stanley, J., … & Klein, R. (2008). Peer Reviewed: Community Health Status Indicators Project: The Development of a National Approach to Community Health. Preventing chronic disease, 5(3).

Newton, R.J., McLellan, S.L., Dila, D.K., Vineis, J.H., Morrison, H.G., Eren, A.M. and Sogin, M.L., 2015. Sewage reflects the microbiomes of human populations. MBio, 6(2), pp.e02574–14.

O’Brien, J.W., Thai, P.K., Eaglesham, G., Ort, C., Scheidegger, A., Carter, S., Lai, F.Y. and Mueller, J.F., (2013). A model to estimate the population contributing to the wastewater using samples collected on census day. Environmental science & technology, 48(1), pp.517–525.

Ort, C., Lawrence, M.G., Rieckermann, J. and Joss, A., (2010). Sampling for pharmaceuticals and personal care products (PPCPs) and illicit drugs in wastewater systems: are your conclusions valid? A critical review. Environmental science & technology, 44(16), pp.6024–6035.

Pehrsson, E.C., Tsukayama, P., Patel, S., Mejía-Bautista, M., Sosa-Soto, G., Navarrete, K.M., Calderon, M., Cabrera, L., Hoyos-Arango, W., Bertoli, M.T. and Berg, D.E., (2016). Interconnected microbiomes and resistomes in low-income human habitats. Nature, 533(7602), p.212.

Piazzon, A., Vrhovsek, U., Masuero, D., Mattivi, F., Mandoj, F., & Nardini, M. (2012). Antioxidant activity of phenolic acids and their metabolites: synthesis and antioxidant properties of the sulfate derivatives of ferulic and caffeic acids and of the acyl glucuronide of ferulic acid. Journal of agricultural and food chemistry, 60(50), 12312–12323.

Pluskal, T., Castillo, S., Villar-Briones, A., & Orešič, M. (2010). MZmine 2: modular framework for processing, visualizing, and analyzing mass spectrometry-based molecular profile data. BMC bioinformatics, 11(1), 395.

Pollet, R.M., D’Agostino, E.H., Walton, W.G., Xu, Y., Little, M.S., Biernat, K.A., Pellock, S.J., Patterson, L.M., Creekmore, B.C., Isenberg, H.N. and Bahethi, R.R., (2017). An atlas of β-glucuronidases in the human intestinal microbiome. Structure, 25(7), pp.967–977.

Preheim, S. P., Perrotta, A. R., Martin-Platero, A. M., Gupta, A., & Alm, E. J. (2013). Distribution-based clustering: using ecology to refine the operational taxonomic unit. Applied and environmental microbiology, AEM-00342.

Prichard, J., Hall, W., de Voogt, P., & Zuccato, E. (2014). Sewage epidemiology and illicit drug research: the development of ethical research guidelines. Science of the Total Environment, 472, 550–555.

Seakins, J.W.T., 1971. The determination of urinary phenylacetylglutamine as phenylacetic acid.: Studies on its origin in normal subjects and children with cystic fibrosis. Clinica Chimica Acta, 35(1), pp.121–131.

Smith, C.A., O’Maille, G., Want, E.J., Qin, C., Trauger, S.A., Brandon, T.R., Custodio, D.E., Abagyan, R. and Siuzdak, G., (2005). METLIN: a metabolite mass spectral database. Therapeutic drug monitoring, 27(6), pp.747–751.

Smith, C. A., Want, E. J., O’Maille, G., Abagyan, R., & Siuzdak, G. (2006). XCMS: processing mass spectrometry data for metabolite profiling using nonlinear peak alignment, matching, and identification. Analytical chemistry, 78(3), 779–787.

Stein, W.H., Paladini, A.C., Hirs, C.H.W. and Moore, S., 1954. Phenylacetylglutamine as a constituent of normal human urine. Journal of the American Chemical Society, 76(10), pp.2848–2849.

Sumner, L.W., Amberg, A., Barrett, D., Beale, M.H., Beger, R., Daykin, C.A., Fan, T.W.M., Fiehn, O., Goodacre, R., Griffin, J.L. and Hankemeier, T., (2007). Proposed minimum reporting standards for chemical analysis. Metabolomics, 3(3), pp.211–221.

Thai, P.K., Jiang, G., Gernjak, W., Yuan, Z., Lai, F.Y. and Mueller, J.F., (2014). Effects of sewer conditions on the degradation of selected illicit drug residues in wastewater. Water research, 48, pp.538–547.

Thomas, K.V., Bijlsma, L., Castiglioni, S., Covaci, A., Emke, E., Grabic, R., Hernández, F., Karolak, S., Kasprzyk-Hordern, B., Lindberg, R.H. and de Alda, M.L., (2012). Comparing illicit drug use in 19 European cities through sewage analysis. Science of the Total Environment, 432, pp.432–439.

United Nations (2018). Revision of the world urbanization prospects. https://population.un.org/wup/

van Nuijs, A. L., Castiglioni, S., Tarcomnicu, I., Postigo, C., de Alda, M. L., Neels, H., … & Covaci, A. (2011). Illicit drug consumption estimations derived from wastewater analysis: a critical review. Science of the Total Environment, 409(19), 3564–3577.

van Nuijs, Alexander LN, Foon Yin Lai, Frederic Been, Maria Jesus Andres-Costa, Leon Barron, Jose Antonio Baz-Lomba, Jean-Daniel Berset et al. (2018). “Multi-year interlaboratory exercises for the analysis of illicit drugs and metabolites in wastewater: development of a quality control system”. TrAC Trends in Analytical Chemistry.

Walley, A. Y., Xuan, Z., Hackman, H. H., Quinn, E., Doe-Simkins, M., Sorensen-Alawad, A., … & Ozonoff, A. (2013). Opioid overdose rates and implementation of overdose education and nasal naloxone distribution in Massachusetts: interrupted time series analysis. Bmj, 346, f174.

Wang, Q., Garrity, G. M., Tiedje, J. M., & Cole, J. R. (2007). Naive Bayesian classifier for rapid assignment of rRNA sequences into the new bacterial taxonomy. Applied and environmental microbiology, 73(16), 5261–5267.

Wishart, D.S., Jewison, T., Guo, A.C., Wilson, M., Knox, C., Liu, Y., Djoumbou, Y., Mandal, R., Aziat, F., Dong, E. and Bouatra, S., (2012). HMDB 3.0—the human metabolome database in 2013. Nucleic acids research, 41(D1), pp.D801–D807.

Wolf, S., Schmidt, S., Müller-Hannemann, M., & Neumann, S. (2010). In silico fragmentation for computer assisted identification of metabolite mass spectra. BMC bioinformatics, 11(1), 148.

Yan, T., O’Brien, P., Shelton, J. M., Whelen, A. C., & Pagaling, E. (2018). Municipal Wastewater as a Microbial Surveillance Platform for Enteric Diseases: A Case Study for Salmonella and Salmonellosis. Environmental science & technology, 52(8), 4869–4877.

Zuccato, E., Chiabrando, C., Castiglioni, S., Bagnati, R. and Fanelli, R., (2008). Estimating community drug abuse by wastewater analysis. Environmental health perspectives, 116(8), p.1027.

