## Supplemental Figures and Tables for "24-hour multi-omics analysis of residential sewage reflects human activity and informs public health"

__Supplementary Figures__

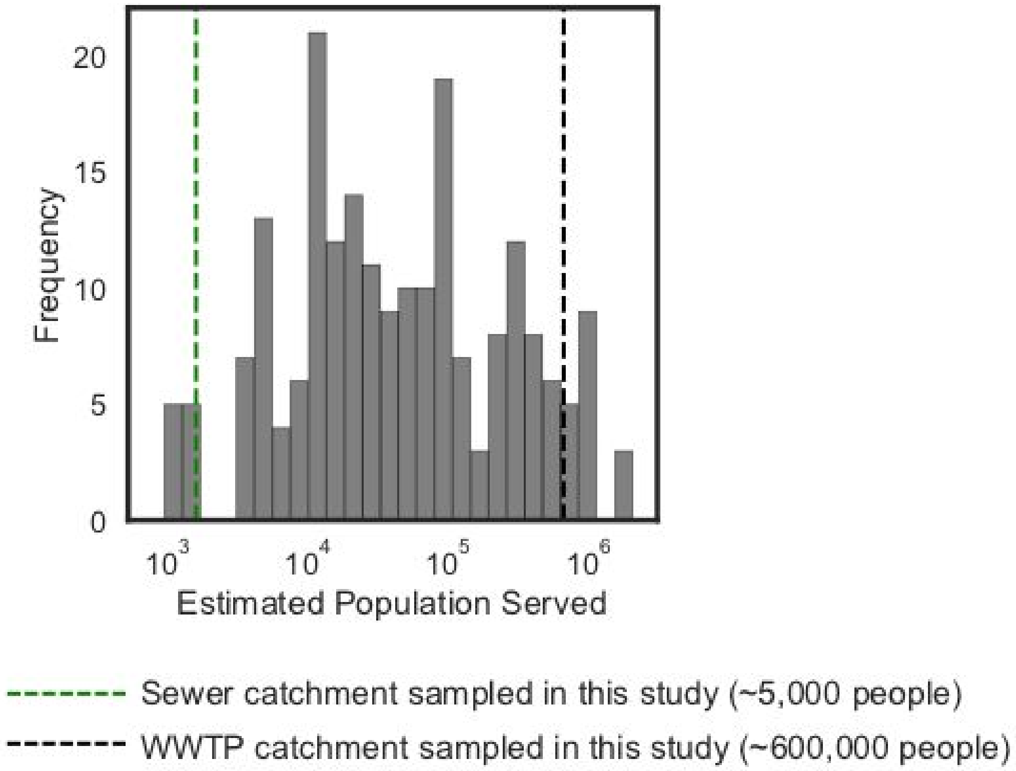


**Supplementary Figure 1.** Distribution of estimated population served by the different WWTP catchments surveyed in Newton et al 2015. Vertical dashed lines indicate the estimated population size of the residential sewer catchment and the WWTP in this study.

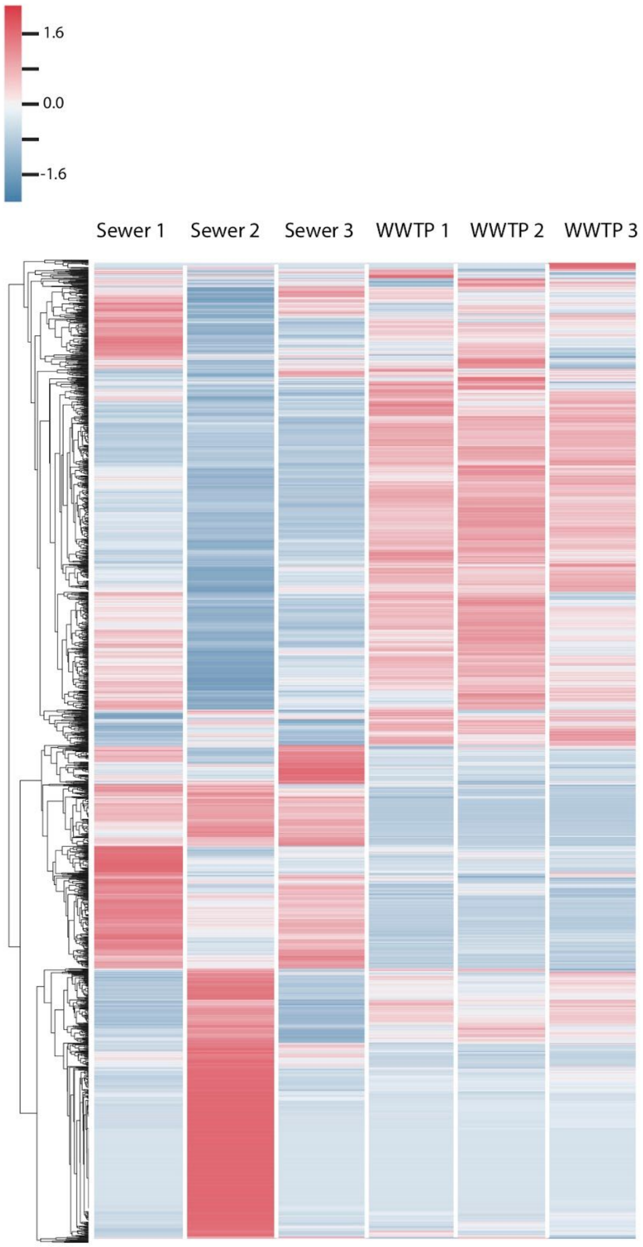


**Supplementary Figure 2.** Heatmap of metabolic features in the residential catchment (n = 3 upstream samples) and at the WWTP catchment (n = 3 downstream samples) clustered by the metabolites' z-scores across all samples. Related to Main Text Figure 2A.

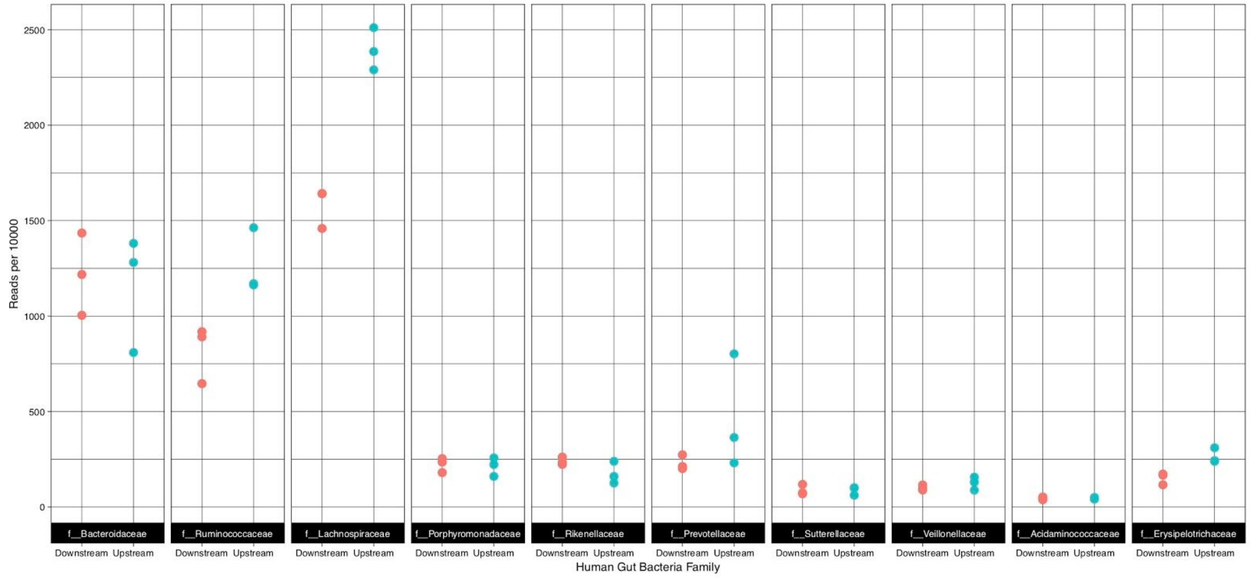


**Supplementary Figure 3.** Relative abundance of the top 10 human fecal bacterial families in WWTP catchment (downstream, red, n = 3) and residential catchment (upstream, blue, n = 3) samples. Related to Main Text Figure 2B.

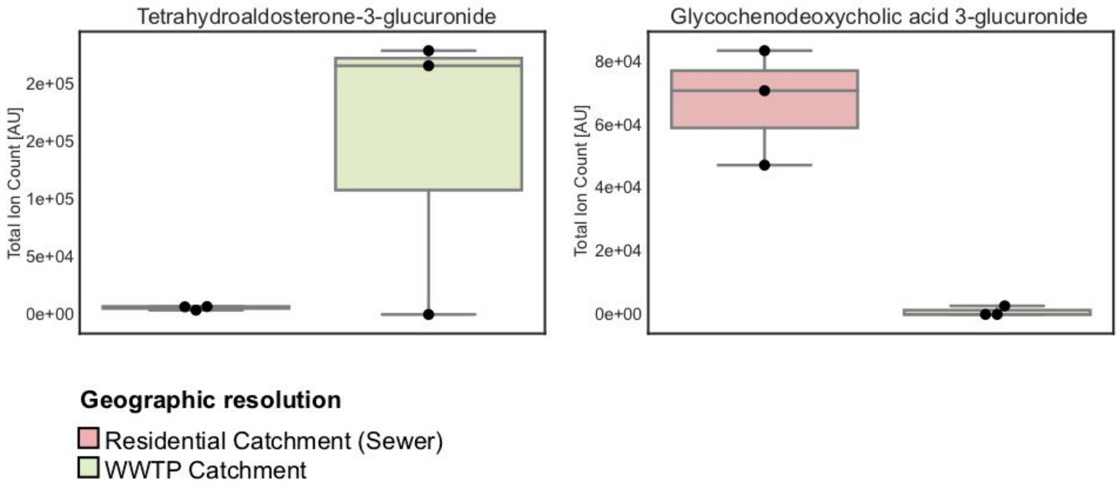


**Supplementary Figure 4.** Additional glucuronides measured in this study and identified at level 2, which are not shown in Figure 2C. Related to Main Text Figure 2C.

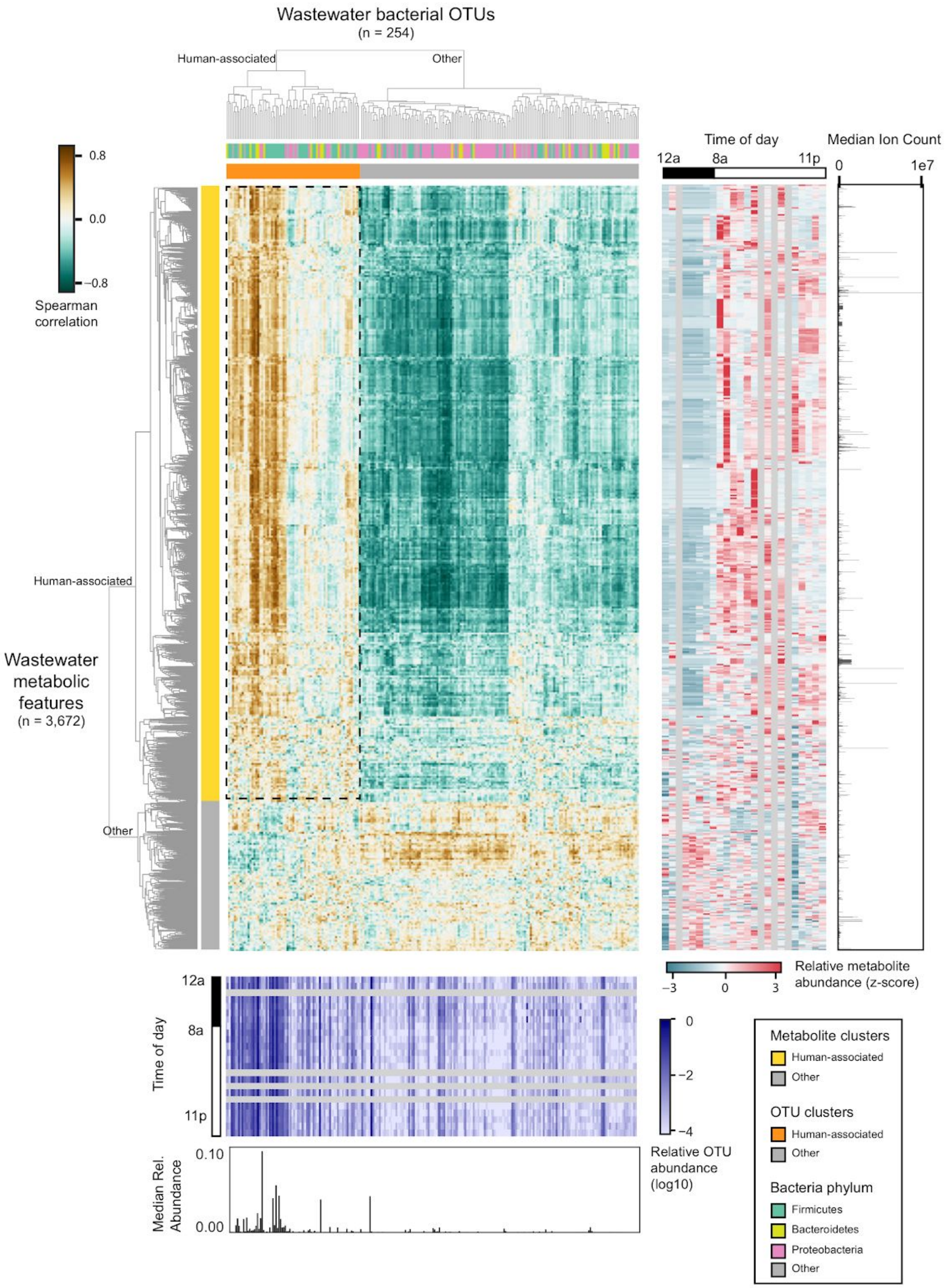


**Supplementary Figure 5.** Co-clustering of metabolites and OTUs. (Top left panel) Heatmap of OTU-metabolite Spearman correlations. Human-associated and non-human-associated features are indicated with the orange and yellow (human-associated) and gray (non-human-associated) bars to the left and top of the heatmap. OTU phyla are indicated on the top panel. (Bottom panel) Top: heatmap of the log10 relative abundance of each OTU (columns) in each time point (rows). Bottom: Median relative abundance of each OTU. (Right panel) Left: heatmap of the relative (z-scored) metabolite abundance for each metabolite (rows) over each time point (columns). Right: median abundance (ion count) of each metabolite.

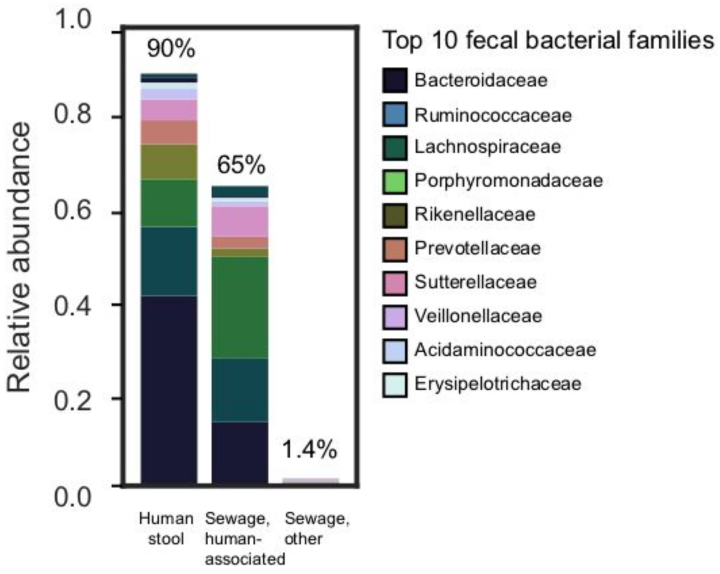


**Supplementary Figure 6.** Relative abundance of the top 10 fecal bacterial communities in human stool from the HMP and the human associated and non-human associated groups in residential catchment sewage. Related to Main Text Figure 3.

### Supplementary Tables

**Supplementary Table 1.** 16S data processing parameters. Custom scripts can be found in https://github.com/thomasgurry/amplicon_sequencing_pipeline and documentation describing the OTU processing pipeline is available at https://amplicon-sequencing-pipeline.readthedocs.io. References: usearch8 (Edgar 2010) and RDP tools (Wang 2007).

| OTU processing step | Function call | Parameter |
|---|---|---|
| Read dereplication | Custom dereplication script | min_count = 10 |
| Length truncation | usearch8 -fastq_filter -fastq_trunclen | 110 |
| Quality trimming | usearch8 -fastq_filter -fastq_truncqual | 25 |
| Similarity-based *de novo* clustering | usearch8 -cluster_otus -otu_radius_pct | 99% |
| RDP taxonomy assignment | RDP classifier (RDPTools/classifier.jar) and custom parsing script | c = 0.5 |

**Supplementary Table 2.** (Excel file, u2-24hr-annotated-metabolites.xlsx) Annotated metabolites.

### Supplementary References

Edgar, R. C. (2010). Search and clustering orders of magnitude faster than BLAST. *Bioinformatics*, *26*(19), 2460-2461.

Newton, R.J., McLellan, S.L., Dila, D.K., Vineis, J.H., Morrison, H.G., Eren, A.M. and Sogin, M.L., 2015. Sewage reflects the microbiomes of human populations. *MBio*, *6*(2), pp.e02574-14.

Wang, Q., Garrity, G. M., Tiedje, J. M., & Cole, J. R. (2007). Naive Bayesian classifier for rapid assignment of rRNA sequences into the new bacterial taxonomy. *Applied and environmental microbiology*, *73*(16), 5261-5267.
